# *GSTM1* copy number is not associated with risk of kidney failure in a large cohort

**DOI:** 10.1101/567651

**Authors:** Yanfei Zhang, Waleed Zafar, Dustin N. Hartzel, Marc S. Williams, Adrienne Tin, Alexander R. Chang, Regeneron Genetics Center, Ming Ta M. Lee, on behalf of the DiscovEHR collaboration

## Abstract

Deletion of *glutathione S-transferase µ1 (GSTM1)* is common in populations and has been asserted to associate with chronic kidney disease progression in some research studies. The association needs to be validated. We estimated *GSTM1* copy number using whole exome sequencing data in the DiscovEHR cohort. Kidney failure was defined as requiring dialysis or receiving kidney transplant using data from the electronic health record and linkage to the United States Renal Data System, or the most recent eGFR < 15 ml/min/1.73m^2^. In a cohort of 46,983 unrelated participants, 28.8% of blacks and 52.1% of whites had 0 copies of *GSTM1*. Over a mean of 9.2 years follow-up, 645 kidney failure events were observed in 46,187 white participants, and 28 in 796 black participants. No significant association was observed between *GSTM1* copy number and kidney failure in Cox regression adjusting for age, sex, BMI, smoking status, genetic principal components, or co-morbid conditions (hypertension, diabetes, heart failure, coronary artery disease, and stroke), whether using a genotypic, dominant, or recessive model. In sensitivity analyses, *GSTM1* copy number was not associated with kidney failure in participants that were 45 years or older at baseline, had baseline eGFR < 60 ml/min per 1.73 m^2^, or with baseline year between 1996-2002. In conclusion, we found no association between *GSTM1* copy number and kidney failure in a large cohort study.

**Translational Statement:** Deletion of *GSTM1* has been shown to be associated with higher risk of kidney failure. However, inconsistent results have been reported. We used electronic health record and whole exome sequencing data of a large cohort from a single healthcare system to evaluate the association between *GSTM1* copy number and risk of kidney failure. We found no significant association between *GSTM1* copy number and risk of kidney failure overall, or in multiple sensitivity and subgroup analyses.

## Introduction

*Glutathione S-transferase µ 1* (*GSTM1*), belongs to the family of glutathione-S-transferases that metabolize a broad range of reactive oxygen species and aldehydes.^1,2^ Loss of *GSTM1* is very common in the population; approximately 50% of whites and 27% of blacks have zero copies of *GSTM1*.^3^ Deletion of one or both copies of the gene results in reduced amount of GSTM1,^4^ and could lead to increased oxidative stress due to diminished ability to neutralize reactive chemical species. An association between loss of *GSTM1* and chronic kidney disease (CKD) progression has been reported in the African American Study of Kidney and Hypertension (AASK).^5^ The association between *GSTM1* copy number and kidney failure was reported in the Atherosclerosis Risk in Communities (ARIC) study which had a larger sample size, both in blacks and whites.^6^ However, data from smaller case-control studies have had mixed findings.^7–11^ The inconsistent results may be due to insufficient sample size and different populations and disease background.

The Geisinger MyCode® Community Health Initiative is an EHR-linked biobank for precision medicine research.^12^ The population served by Geisinger have low rates of out migration, thus the electronic health record (EHR) data are relatively complete longitudinally, with a median of 14 years of follow-up.^12^ Also, a high proportion of participants were found to have first and second-degree relatedness through cryptic relatedness analysis.^13^ Through the ongoing DiscovEHR collaboration with the Regeneron Genetic Center, whole exome sequence (WES) data are available from approximately 92,000 MyCode® participants to date.^12,14,15^ These comprehensive clinical data are linked to matched genetic data and provide power to identify disease-genetic associations.^16^

Confirming whether or not loss of *GSTM1* increases the risk of kidney failure is important, given the high prevalence of *GSTM1* loss in the population and the serious morbidity and mortality associated with kidney failure. In this study, we examine the relationship between *GSTM1* copy number and incident kidney failure in the Geisinger MyCode cohort.

## Results

### Study cohort and baseline characteristics

There were 46,983 unrelated participants included in the main analysis. Among these, 46,187 (98.3%) were white, and 796 (1.7%) were black. Frequency of *GSTM1* copy number differed between blacks and whites (Supplemental Figure 4). Almost half of the blacks (49.2%) had 1 copy of *GSTM1* compared with only 39.6% of whites, and more than half (52.1%) of whites had 0 copies of *GSTM1* compared with 28.8% of blacks. Chi-square test indicated *GSTM1* copy number follows the HWE (Table 1. P-values were 1*10^-4^ for whites and 0.618 for blacks).

**Table 1:** Baseline characteristics of participants by *GSTM1* copy number

| Characteristic | Whites |  |  |  | Blacks |  |  |  |
| --- | --- | --- | --- | --- | --- | --- | --- | --- |
| Copy number | 0 | 1 | 2 | P | 0 | 1 | 2 | P |
| N. | 24141 | 18255 | 3791 | 1E-4 ^ | 233 | 388 | 175 | 0.618 ^ |
| Age | 50.9(14.1) | 50.8(14.0) | 51.2(14.3) | 0.317 | 41.8(12.3) | 44.7(12.1) | 44.3(13.2) | 0.015 |
| Female, n(%) | 14184(58.8) | 10750(58.9) | 2208(58.2) | 0.763 | 148(63.5) | 240(61.9) | 103(58.9) | 0.629 |
| eGFR | 91.8(20.3) | 91.7(20.4) | 91.3(20.8) | 0.326 | 95.0(22.9) | 90.8(23.2) | 91.9(24.4) | 0.095 |
| BMI | 31.6(7.9) | 31.6(7.8) | 31.6(7.9) | 0.817 | 33.5(9.3) | 34.7(9.6) | 34.5(10.2) | 0.299 |
| Smoking Status |  |  |  |  |  |  |  |  |
| Current Smoker | 4688 (19.4) | 3521 (19.3) | 769 (20.3) | 0.418 | 65 (27.9) | 109 (28.1) | 55 (31.4) | 0.785 |
| Former Smoker | 7803 (32.3) | 5829 (31.9) | 1231 (32.5) |  | 64 (27.5) | 117 (30.2) | 52 (29.7) |  |
| Never Smoker | 11650 (48.3) | 8905 (48.8) | 1791 (47.2) |  | 104 (44.6) | 162 (41.8) | 68 (38.9) |  |
| Hypertension | 7193 (29.8) | 5409 (29.6) | 1118 (28.4) | 0.893 | 96 (41.2) | 169 (43.6) | 83 (47.4) | 0.453 |
| T1DM | 251 (1.0) | 205 (1.1) | 50 (1.3) | 0.277 | 6 (2.5) | 12 (3.1) | 5 (2.9) | 0.932 |
| T2DM | 3130 (13.0) | 2299 (12.6) | 483 (12.7) | 0.522 | 40 (17.2) | 92 (23.7) | 35 (20.0) | 0.143 |
| CAD | 1384 (5.7) | 1072 (5.9) | 241 (6.4) | 0.304 | 0 | 0 | 0 | / |
| HF | 568 (2.4) | 402 (2.2) | 110 (2.9) | 0.034 | 10 (4.3) | 12 (3.1) | 7 (4.0) | 0.713 |
| Stroke | 413(1.7) | 304(1.7) | 69(1.8) | 0.789 | 4(1.7) | 11(2.8) | 8(4.6) | 0.233 |

^ Hardy-Weinberg equilibrium test using Chi-Square test. Consider the sample size for Whites (46,187), we selected a more stringent significance level of 1E-5.
P<0.0045 (0.05/11) was considered statistically significant. T1DM: Type 1 diabetes mellitus. T2DM: type 2 diabetes mellitus. CAD: coronary artery disease; HF: heart failure.

Baseline demographic and clinical characteristics of the participants stratified by race and *GSTM1* copy numbers are provided in table 1. The average baseline age for whites and blacks was 51 and 44 years old respectively. The mean baseline eGFR were 91 and 92 ml/min per 1.73 m^2^. Prevalence of hypertension was 29% in whites and 44% in blacks, and prevalence of type 2 diabetes was 13% in whites and 21% in blacks at baseline. No statistically significant differences were observed for baseline characteristics among the three genotype groups when adjusted for multiple comparisons.

### Time-to-event analysis

Over a mean follow-up of 9.2 years, there were a total of 645 kidney failure events in 46,187 white participants. Over a mean follow-up of 5.5 years, there were 28 events in 796 blacks. No significant difference in kidney failure-free survival was found across *GSTM1* copy numbers by log rank test (P=0.9 and 0.5 for whites and blacks, respectively, Figure 1).

**Figure 1:**
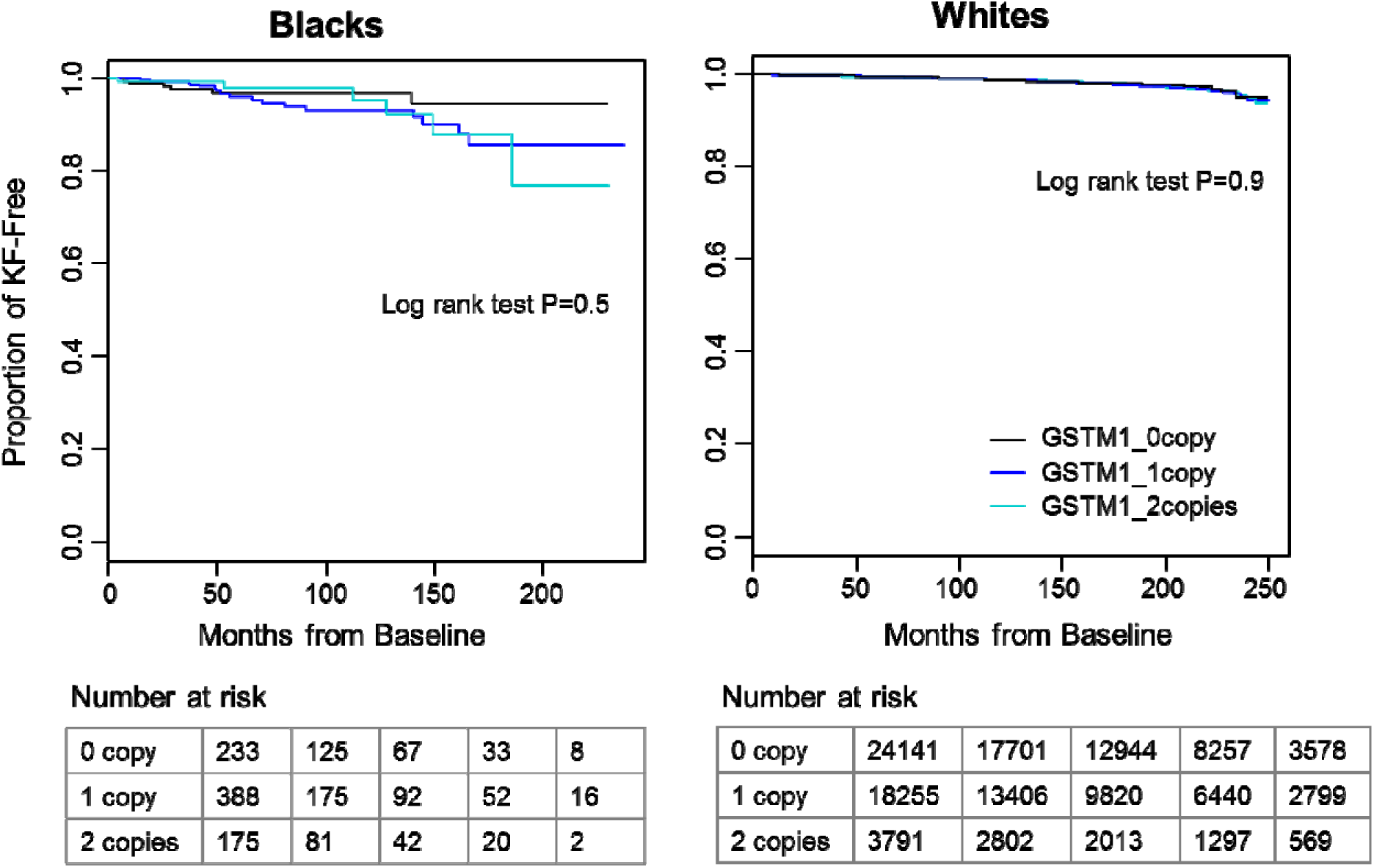
Kaplan-Meier survival curves of time to kidney failure by *GSTM1* copy number in white and black participants. In both races, participants without *GSTM1* (black line) had longer kidney failure-free survival time than those with 1 or 2 copies of *GSTM1* (blue and cyan lines). No significant difference was found by log rank test. Tables below the figure showed the number of patients at risk with start time at Month 0, 50, 100, 150 and 200.

Findings were similar in Cox regression analyses adjusting for other covariates (Table 2). In the genotypic model, after adjusting for age, sex and the first 4 PCs, there was no difference in risk of kidney failure among white participants with 0 copies (hazard ratio [HR] 1.01, 95% confidence interval [CI]: 0.81-1.48]) or 1 copy (HR 1.08, 95% CI: 0.80-1.47), compared to those with 2 copies. Findings were similar for black participants in the genotypic model adjusting for age, sex, and the first 4 PCs (0 copies HR 0.68, 95% CI: 0.22-2.15; 1 copy HR 1.13, 95% CI: 0.44-2.90). In the dominant model, after adjusting for age, sex, and the first 4 PCs, there was again no difference in risk of kidney failure among participants with 0 or 1 copies for white participants (HR 1.10, 95% CI: 0.82-1.46) or black participants with 0 or 1 copies (HR 0.96, 95% CI: 0.39-2.34), compared to their counterparts with 2 copies. In the recessive model, after adjusting for age, sex, and the first 4 PCs, there was no difference in risk of kidney failure among white participants with 0 copies (1.03, 95% CI: 0.88-1.20), or black participants with 0 copies (HR 0.63, 95% CI: 0.25-1.56), compared to their counterparts with 1 or 2 copies. Adjustment for additional renal risk factors in Model 2 yielded similar results (Table 2 and Supplemental Tables 1-3).

**Table 2:** Risk of kidney failure associated with *GSTM1* copy number

| Population | Hazard Ratio [95% CI] |  |  |  |  |  |  |  |
| --- | --- | --- | --- | --- | --- | --- | --- | --- |
|  | Genotypic Model (Ref: 2 copies) |  |  |  | Dominant Model |  | Recessive Model |  |
|  | 0 copy | P | 1 copy | P | 0/1 vs 2 | P | 0 vs 1/2 | P |
| Blacks (28/796) |  |  |  |  |  |  |  |  |
| Model 1 | 0.68 [0.22, 2.15] | 0.516 | 1.13 [0.44, 2.90] | 0.802 | 0.96 [0.39, 2.34] | 0.937 | 0.63 [0.25, 1.56] | 0.316 |
| Model 2 | 0.67 [0.19, 2.34] | 0.527 | 0.63 [0.22, 1.84] | 0.399 | 0.64 [0.23, 1.77] | 0.391 | 0.93 [0.33, 2.59] | 0.883 |
| Whites (645/46187) |  |  |  |  |  |  |  |  |
| Model 1 | 1.01 [0.81, 1.48] | 0.537 | 1.08 [0.80, 1.47] | 0.604 | 1.10 [0.82, 1.46] | 0.551 | 1.03 [0.88, 1.20] | 0.732 |
| Model 2 | 1.14 [0.84, 1.54] | 0.398 | 1.15 [0.84, 1.56] | 0.384 | 1.14 [0.85, 1.53] | 0.375 | 1.02 [0.87, 1.19] | 0.836 |
| Whites, Baseline Age≥45 (518/31406) |  |  |  |  |  |  |  |  |
| Model 1 | 1.03 [0.74, 1.42] | 0.878 | 0.99 [0.71, 1.37] | 0.931 | 1.01 [0.74, 1.34] | 0.960 | 1.04 [0.87, 1.23] | 0.673 |
| Model 2 | 1.12 [0.81, 1.54] | 0.504 | 1.07 [0.77, 1.49] | 0.677 | 1.10 [0.80, 1.50] | 0.562 | 1.05 [0.89, 1.25] | 0.553 |
| Whites, Baseline eGFR<60 (304/3129) |  |  |  |  |  |  |  |  |
| Model 1 | 1.13 [0.75, 1.69] | 0.556 | 1.04 [0.69, 1.57] | 0.860 | 1.09 [0.74, 1.61] | 0.670 | 1.10 [0.87, 1.37] | 0.429 |
| Model 2 | 1.03 [0.68, 1.55] | 0.891 | 1.03 [0.68, 1.56] | 0.897 | 1.03 [0.69, 1.53] | 0.889 | 1.01 [0.80, 1.27] | 0.956 |
| Whites, Baseline Year before 2003 (310/14572) |  |  |  |  |  |  |  |  |
| Model 1 | 0.88 [0.59, 1.32] | 0.542 | 0.88 [0.59, 1.33] | 0.553 | 0.88 [0.60, 1.30] | 0.529 | 0.98 [0.78, 1.22] | 0.841 |
| Model 2 | 0.86 [0.57, 1.29] | 0.460 | 0.89 [0.59, 1.34] | 0.563 | 0.87 [0.59, 1.29] | 0.485 | 0.95 [0.76, 1.19] | 0.648 |

Model 1: adjusted for age, sex, first 4 genetic principle components. Model 2: Model 1 plus baseline eGFR, BMI, smoking status, hypertension, type 1 diabetes, type 2 diabetes, heart failure, coronary artery disease, and stroke.

### Sensitivity analyses

Analyses were performed on white participants who were older than 45 years of age at baseline (n=31406), had baseline eGFR < 60 ml/min/1.73m^2^ (n=3129), or had index date before 2003 (n=14572). No statistically significant differences in the risk of kidney failure were identified for participants older than 45 years of age at baseline (0 copies: HR 1.03. 95% CI: 0.74-1.42; 1 copy: HR 0.99. 95% CI: 0.71-1.37), or those with baseline eGFR < 60 ml/min/1.73m^2^ (0 copies: HR 1.13, 95% CI: 0.75-1.69; 1 copy HR 1.04, 95% CI: 0.69-1.57), or for those with baseline year before 2003 (0 copies: HR 0.88, 95% CI: 0.59-1.32; 1 copy HR 0.88. 95% CI: 0.59-1.33) (Table 2 and Supplemental Tables 1-3).

## Discussion

To our knowledge, this is the largest study to investigate the association of *GSTM1* copy number variation with kidney failure. There were a total of 46,187 unrelated whites and 796 blacks in the study with an average of 9.3 years’ follow-up. The frequency of the *GSTM1*copy numbers were very similar to those reported previously.^3,5,6^ Our results showed no significant association between *GSTM1* copy number and risk of kidney failure in unadjusted or adjusted analyses whether using a genotypic, dominant, or recessive genetic model.

Data from the ARIC and AASK cohorts suggested that the loss of *GSTM1* increased the risk of kidney failure or accelerated CKD progression.^5,6,17^ In ARIC, a community-based cohort of middle-aged black and white participants, there were 3461 white participants with WES reads. In fully adjusted models among whites in ARIC, loss of *GSTM1* was associated with risk of kidney failure (0 or 1 copy vs. 2 copies: HR 2.54; 95% CI: 1.32-4.88). It is possible that differences in baseline characteristics of the study populations could explain differing findings. Our cohort had a lower prevalence of smoking (never smoking 48% vs. 38%), a higher prevalence of diabetes (14% vs. 8%), and was more contemporary (median baseline year 2004 vs. ARIC baseline year 1987-1989). However, in sensitivity analyses in 14572 white participants with baseline year between 1996-2002 in our cohort, there was no association between *GSTM1* loss and risk of kidney failure.

There is also the possibility that *GSTM1* may only be deleterious when GFR falls below certain levels, or in the setting of specific types of kidney disease. In the AASK study, a randomized trial comparing different antihypertensive medications and levels of blood pressure control in blacks with CKD attributed to hypertension, 692 participants had *GSTM1* genotyping completed. Loss of GSTM1 in AASK was associated with increased risk of CKD progression (HR 0 copy: HR 1.88; 95% CI: 1.07-3.30; HR 1 copy: 1.68; 95% CI: 1.00-2.84). While there were few blacks in our study population with CKD, we found no association between loss of *GSTM1* and risk of kidney failure in 3129 white participants with baseline eGFR < 60 ml/min/1.73m^2^. Some smaller case-control studies also tested a recessive genetic model (0 copies vs. 1 or 2 copies) of *GSTM1* and found that *GSTM1* 0 copy was associated with kidney failure in several case-control studies.^5,10,11,18^ In our study, however, we examined both recessive and dominant genetic models, and found no associations between *GSTM1* copy number and risk of kidney failure. Some limitations of this study are worth noting. First, the genotypes of *GSTM1* in this study were derived from WES and the thresholds were determined empirically from the histogram. Multiplex PCR validation was not performed for this study. However, this method was used in the ARIC study, which showed 99.3% agreement when comparing the results of 0 copy with ≥ 1 copy from PCR.^6^ Moreover, the frequency of *GSTM1* copy numbers in this study is similar to those reported previously for both whites and blacks,^5,6^ and it follows the Hardy-Weinberg equilibrium. We also examined the possibility of miss-mapping due to the complexity of *GSTM* locus. The histogram of the sum of normalized coverage of *GSTM5*, the paralog gene of *GSTM1*, is unimodal distribution. Based on these evidences, we believe the copy number results derived from WES had a low error rate and can be used with confidence for analysis. Second, the sample size for blacks was small with only 796 patients available for analysis, limiting the power to examine an association between *GSTM1* and kidney failure in this population. The strength of this study is the large sample size of whites, which included 46,187 patients, allowing for large subgroup analyses in older patients, those with baseline eGFR < 60 ml/min/1.73m^2^, and those with longer duration of follow-up. Finally, *GSTM1* deletion is known to be protective for lung cancer among individuals with a high intake of cruciferous vegetable and deleterious among those with low intake of cruciferous vegetable.^19–23^ We were not able to analyze the effect modification between *GSTM1* copy number and cruciferous vegetable intake.

In conclusion, our study does not support an association between loss of *GSTM1* and increased risk of kidney failure. Additional research is needed to confirm whether loss of *GSTM1* increases risk of kidney failure in certain subgroups such as blacks, and the interaction with diet.

## Methods

### Study Population

The study population included 72,756 participants in the Geisinger-Regeneron DiscovEHR cohort with WES data and at least 3 serum creatinine values. We excluded participants who had baseline eGFR < 15 ml/min/1.73m^2^ or history of dialysis or transplant (n=417), baseline age <18 (n=1,960), missing BMI values (n=230), and unknown smoking status (n=2,694). Cryptic relatedness was evaluated using IBD method in PLINK1.9.^24^ One of the related pair of participants with PI_HAT >= 0.125 were removed to reduce confounding from other shared genetic/environmental factors that could not be assessed, resulting in 46983 participants (Supplement Figure 1). We estimated eGFR using the CKD-Epidemiology Collaboration equation.^25,26^ We defined the study period, from a baseline time of the second serum creatinine to the time of a renal endpoint, or the last serum creatinine test. All participants provided their informed written consent, and the study was approved by the Geisinger Institutional Review Board.

### Outcome definition

An assignment of kidney failure was made if any of the following criteria was met: 1) the last available eGFR was less than 15 mL/min/1.73 m^2^; 2) the EHR showed an International Classification of Disease (ICD) code for end stage renal disease (ESRD) (ICD9: 585.6, ICD10: N18.6); 3) receipt of dialysis or transplant per linkage to the United States Renal Data System (USRDS). For participants who met more than one criteria, the earliest documented date of the criterion was considered the kidney failure date.

### Clinical variables

The following data elements were extracted from the EHR: baseline age, sex, self-reported race, body mass index (BMI, BMI = Weight(kg)/Height(m)^2^), serum creatinine, smoking status; and ICD-9/10 coded diagnosis of hypertension, diabetes, coronary artery disease, heart failure, and stroke. Genetic principal components were calculated by PLINK1.9 (www.cog-genomics.org/plink/1.9/).^24^

### *GSTM1* and *GSTM5* copy number estimation

DNA was sequenced in two batches at the Regeneron Genetic Center. The WES data processing has been described elsewhere,^13,15^ and details are also provided in the supplementary methods. Copy number of *GSTM1* was estimated using the method reported previously.^6^ Briefly, sequence coverage was normalized using CLAMMS software.^27^ Normalized coverage of all eight exons of *GSTM1* was summed for each participant. Thresholds for *GSTM1* copy number were determined empirically according to the distribution of sum of normalized coverage (Supplementary Figure 2). The copy number of *GSTM5* was estimated the same way as *GSTM1* (Supplementary notes).

### Statistical analyses

Hardy-Weinberg equilibrium (HWE) was estimated using Chi-Square test for *GSTM1* copy number. A more stringent significance level of 1e^-5^ was selected due to the large sample size (46,187) for Whites. Nominal significance level of 0.05 was used for Blacks as the small sample size (796). Missing baseline BMI values were imputed using mean BMI values calculated from all available BMI values for each participant, and missing smoking status at baseline was imputed using the most recent recorded smoking status in the EHR (Supplemental Figure 3). Baseline characteristics were compared using *ANOVA* for continuous variables, and chi-squared tests for categorical variables. A p-value of < 0.05/*N* was considered statistically significant after Bonferroni adjustment for multiple comparisons, where *N* is the number of variables compared. Kaplan–Meier curves were plotted by *GSTM1* copy number, and survival differences between genotype groups were assessed using a log rank test. All subsequent analyses were stratified by race given the difference in allele frequency and sample size. Two models were evaluated. In Model 1, Cox proportional hazards model was adjusted for age, sex and the first four genetic principle components; Based on Model 1, Model 2 was additionally adjusted for risk factors including baseline eGFR, smoking status, BMI, hypertension, diabetes, coronary artery disease, heart failure and stroke. Three genetic models were evaluated: 1) genotypic model, using copy number 0,1, and 2 as three categories; 2) dominant model (0 or 1 copy vs. 2 copies *GSTM1)*; 3) recessive model (1 or 2 copies vs. 0 copies of *GSTM1).* To explore whether the effect of *GSTM1* loss was stronger in specific higher-risk subgroups, sensitivity analyses were completed including subsets of 1) older participants (baseline age ≥ 45 years); 2) participants with CKD (baseline eGFR < 60 ml/min per 1.73 m^2^); 3) participants with longer follow-up (baseline year 1996-2002). Power was estimated using powerCT() function in R package powerSurvEpi 0.1.0. Given the current sample size and event number, we have power of >=0.8 to test hazard ratio of at least 1.4 and 2.8 for white and black cohort, respectively. All analyses were performed using R (version 3.4.3).

## Disclosure

Regeneron Genetics Center authors work for Regeneron Pharmaceuticals.

## Acknowledgments

The authors thank the staff and participants of MyCode. The Regeneron Genetics Center funded the collection of study samples, the generation of whole exome sequencing data. Geisinger provided funding for clinical data extraction and other analysis. A.C. is supported by National Institutes of Health/National Institute of Diabetes and Digestive and Kidney Diseases grant K23 DK106515-01. Data reported here were supplied by the U.S. Renal Data System. The interpretation and reporting of these data are the responsibility of the authors and in no way should be seen as an official policy or interpretation of the U.S. government. The authors thank Ilene G. Ladd who is the project manager at Genomic Medicine Institute for English editing.

## Author Contributions

A.T, A.C, and M.T.M.L designed the study; D.N.H extracted data; Y.Z and W.Z performed the analysis and drafted the manuscript. M.S.W, M.T.M.L and A.C did critical review on the paper. RGC supported the study and did review the manuscript. All authors approved the final version of the manuscript.

