## Supplementary Notes and Figures for "*GSTM1* copy number is not associated with risk of kidney failure in a large cohort"

The sources of support that require acknowledgment: The Regeneron Genetics Center funded the collection of study samples, the generation of whole exome sequencing data.

**Running headline**: *GSTM1* and risk of kidney failure

### **Supplementary methods:**

#### **Sample preparation and whole exome sequence data processing**

Sample prep and sequencing for the first ~61K samples have been previously described ^1^ and this set of samples are referred to in this supplementary note as the “60k”. The additional set of ~31K samples were prepared in the same process and was described recently;^2^ raw sequencing data was gathered in local buffer storage and uploaded to the DNAnexus (Mountain View, CA) platform for automated analysis. Sample-level read files were generated with CASAVA (Illumina Inc.) and aligned to GRCh38 with BWA-mem.^3^ The resultant BAM files were processed using GATK (<https://software.broadinstitute.org/gatk/>) ^4^ and Picard (<http://broadinstitute.github.io/picard/> ) to sort, mark duplicates, and perform local realignment of reads around putative indels.

#### **Coverage Normalization using CLAMMS**

CLAMMS ^5^ utilizes each sample’s BAM file, specifically using base-level depth-of-coverage calculations from aligned reads having mapping quality >= 30, and seven sequencing quality control metrics computed using Picard. Depth-of-coverage profiles are generated for exon “windows” representing either the entire exon or a 500-1000 base pair contiguous segment of an exon for exons that span more than 1000 base pairs in length. Exon window coverage distributions are normalized independently for every sample, adjusting for GC content and overall sequencing depth.

#### ***GSTM1* and *GSTM5* Coverage Distribution**

The normalized coverage for the eight exons of *GSTM1* were then extracted and summed for each individual. The *GSTM1* coordinates used were chr1:109687870-109693299 for 60k and chr1:109687870-109693745 for 30k due to different capture kits. Sum of normalized coverage were then plotted (supplementary figure 2).

The normalized coverage for the eight exons of *GSTM5* were extracted and summed for each individual the same way as *GSTM1*. The *GSTM5* coordinates used were chr1:109712242 to 109718268. Sum of normalized coverage were plotted in supplementary figure 5.

### **Supplementary figures and legends:**

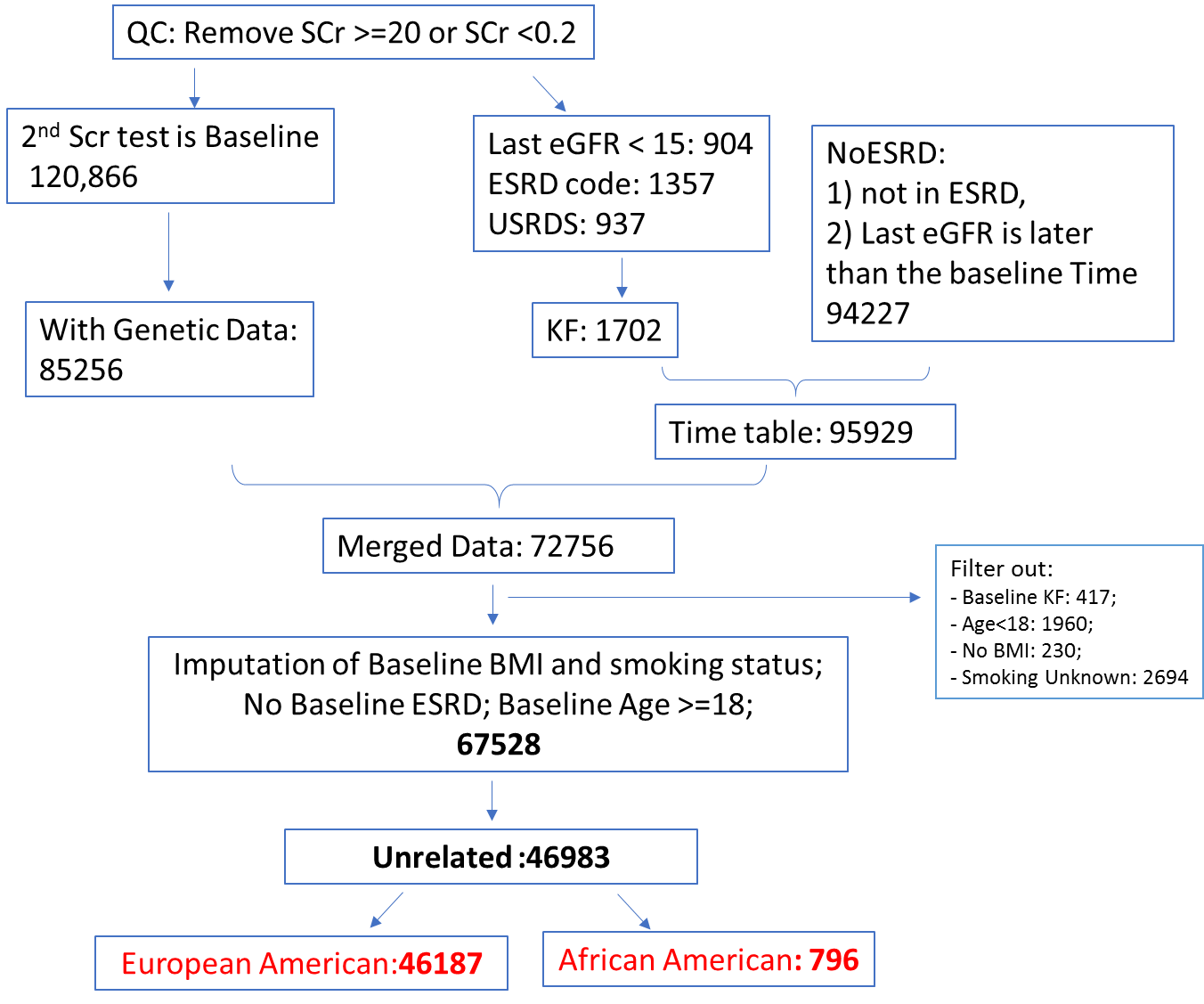

#### **Supplementary figure 1**:

Data processing workflow. Extreme values were removed (SCr >=20 or <0.2). The 2^nd^ SCr test was defined as baseline. Kidney failure was defined as any of the 3 criteria was met: 1) Last eGFR < 15; 2) having ICD codes for ESRD; 3) Registered at USRDS. Participants without genetic data were removed. Participants who had kidney failure or age <18 at baseline were removed. Participants who had unknown smoking status or never had BMI measured were also removed. Cryptic relatedness was calculated by IBD method. One of each pair of the related sample was removed. Finally, 46983 unrelated participants, including 46187 whites and 796 blacks were included in the analysis. SCr: serum creatinine; KF: kidney failure.

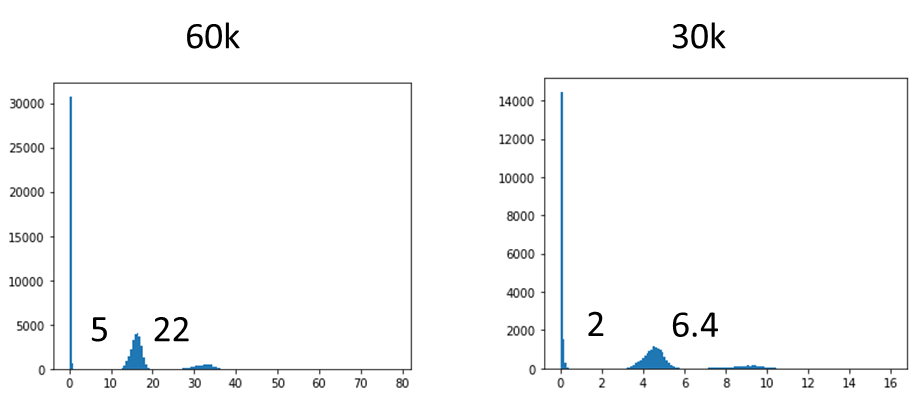

#### **Supplementary figure 2**:

Histogram of sum of normalized average coverage for *GSTM1*. Coverage was normalized using CLAMMS algorithm. Normalized coverage for the eight exomes of *GSTM1* were summed. Cut off value for 0, 1 and 2 copies are set to 5, 22 for 60k and 2, 6.4 for the additional 30k.

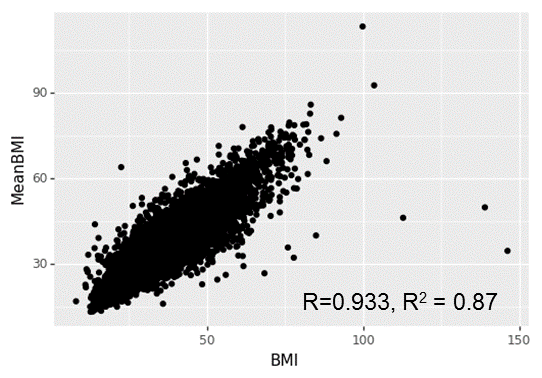

#### **Supplementary figure 3**:

Scatter plot of Mean BMI (Y-axis) and Baseline BMI (X-axis) from 59752 participants. Correlation coefficient was calculated. Mean BMI is highly correlated to the baseline BMI (R^2^ = 0.87).

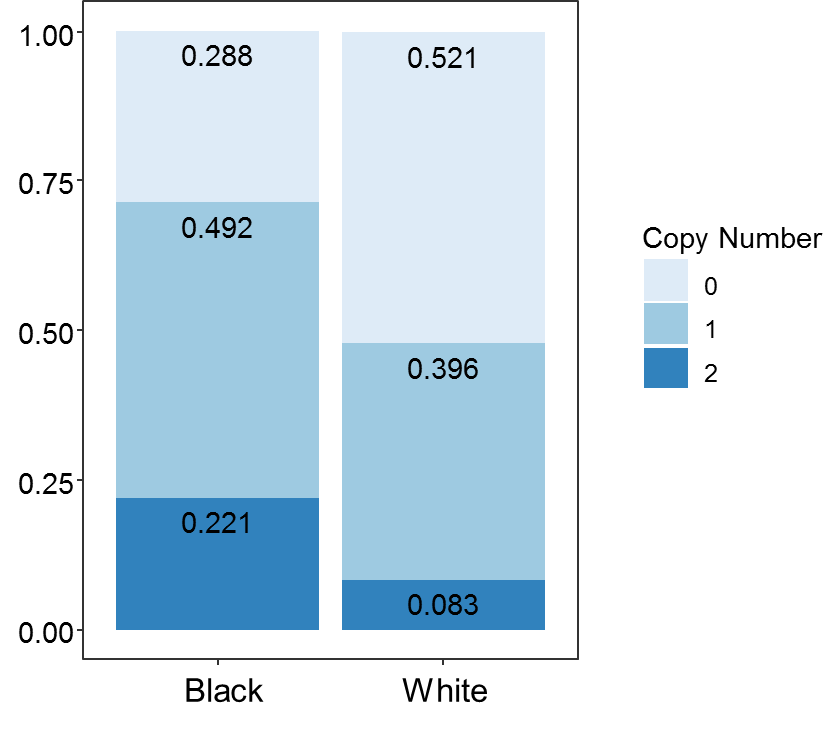

#### **Supplementary figure 4**:

The frequency of *GSTM1* copy numbers in unrelated participants.

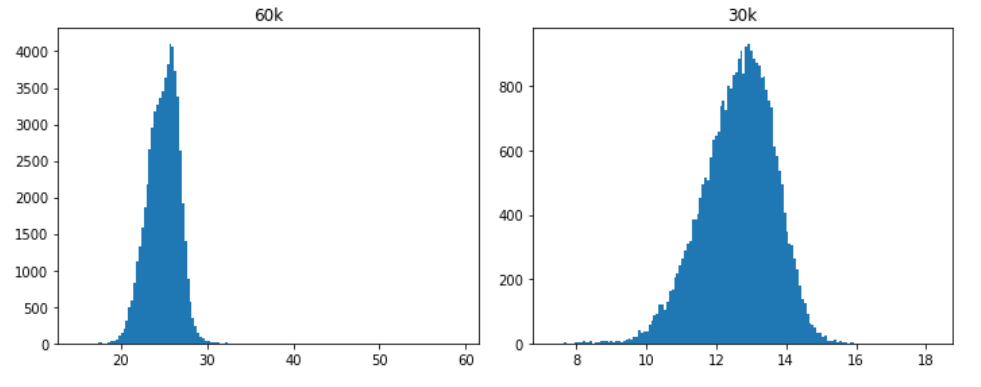

#### **Supplementary figure 5:**

Histogram of sum of normalized average coverage for *GSTM5* in 60k and 30k.

**Regeneron Genetics Center Banner Author List and Contribution Statements**

All authors are listed in alphabetical order.

RGC Management and Leadership Team

Goncalo Abecasis, Ph.D., Aris Baras, M.D., Michael Cantor, M.D., Giovanni Coppola, M.D., Aris Economides, Ph.D., John D. Overton, Ph.D., Jeffrey G. Reid, Ph.D., Alan Shuldiner, M.D.

Contribution: All authors contributed to securing funding, study design and oversight, and review and interpretation of data and results. All authors reviewed and contributed to the final version of the manuscript.

Sequencing and Lab Operations

Christina Beechert, Caitlin Forsythe, M.S., Erin D. Fuller, Zhenhua Gu, M.S., Michael Lattari, Alexander Lopez, M.S., John D. Overton, Ph.D., Thomas D. Schleicher, M.S., Maria Sotiropoulos Padilla, M.S., Karina Toledo, Louis Widom, Sarah E. Wolf, M.S., Manasi Pradhan, M.S., Kia Manoochehri, Ricardo H. Ulloa.

Contribution: C.B., C.F., K.T., A.L., and J.D.O. performed and are responsible for sample genotyping. C.B, C.F., E.D.F., M.L., M.S.P., K.T., L.W., S.E.W., A.L., and J.D.O. performed and are responsible for exome sequencing. T.D.S., Z.G., A.L., and J.D.O. conceived and are responsible for laboratory automation. M.P., K.M., R.U., and J.D.O are responsible for sample tracking and the library information management system.

Genome Informatics

Xiaodong Bai, Ph.D., Suganthi Balasubramanian, Ph.D., Leland Barnard, Ph.D., Andrew Blumenfeld, Yating Chai, Ph.D., Gisu Eom, Lukas Habegger, Ph.D., Young Hahn, Alicia Hawes, B.S., Shareef Khalid, Jeffrey G. Reid, Ph.D., Evan K. Maxwell, Ph.D., John Penn, M.S., Jeffrey C. Staples, Ph.D., Ashish Yadav, M.S.

Contribution: X.B., A.H., Y.C., J.P., and J.G.R. performed and are responsible for analysis needed to produce exome and genotype data. G.E., Y.H., and J.G.R. provided compute infrastructure development and operational support. S.K., S.B., and J.G.R. provide variant and gene annotations and their functional interpretation of variants. E.M., L.B., J.S., A.B., A.Y., L.H., J.G.R. conceived and are responsible for creating, developing, and deploying analysis platforms and computational methods for analyzing genomic data.

Clinical Informatics

Nilanjana Banerjee, Ph.D., Michael Cantor, M.D.

Contribution: All authors contributed to the development and validation of clinical phenotypes used to identify study subjects and (when applicable) controls.

Analytical Genomics and Data Science

Goncalo Abecasis, Ph.D., Amy Damask, Ph.D., Lauren Gurski, Alexander Li, Ph.D., Nan Lin, Ph.D., Daren Liu, Jonathan Marchini Ph.D., Anthony Marcketta, Shane McCarthy, Ph.D., Colm O’Dushlaine, Ph.D., Charles Paulding, Ph.D., Claudia Schurmann, Ph.D., Dylan Sun, Tanya Teslovich, Ph.D., Cristopher Van Hout, Ph.D., Bin Ye

Contribution: Development of statistical analysis plans. QC of genotype and phenotype files and generation of analysis ready datasets. Development of statistical genetics pipelines and tools and use thereof in generation of the association results. QC, review and interpretation of result. Generation and formatting of results for manuscript figures. Contributions to the final version of the manuscript.

Therapeutic Area Genetics

Jan Freudenberg, M.D., Nehal Gosalia, Ph.D., Claudia Gonzaga-Jauregui, Ph.D., Julie Horowitz, Ph.D., Kavita Praveen, Ph.D.

Contribution: Development of study design and analysis plans. Development and QC of phenotype definitions. QC, review, and interpretation of association results. Contributions to the final version of the manuscript.

Planning, Strategy, and Operations

Paloma M. Guzzardo, Ph.D., Marcus B. Jones, Ph.D., Lyndon J. Mitnaul, Ph.D.

Contribution: All authors contributed to the management and coordination of all research activities, planning and execution. All authors managed the review of data and results for the manuscript. All authors contributed to the review process for the final version of the manuscript.
